# Delineating sex-dependent and anatomic decline of motor functions in the SOD1G93A mouse model of amyotrophic lateral sclerosis

**DOI:** 10.1101/2024.12.17.628968

**Authors:** Oksana Shelest, Ian Tindel, Marie Lauzon, Ashley Dawson, Ritchie Ho

**Author notes:** Correspondence: Ritchie Ho. These authors contributed equally to this work (Oksana Shelest, Ian Tindel).

## Abstract

The transgenic SOD1G93A mouse model is the most widely used animal model of amyotrophic lateral sclerosis (ALS), a fatal disease of motor neuron degeneration. While genetic background influences onset and progression variability of motor dysfunction, the C57BL/6 background most reliably exhibits robust ALS phenotypes; thus, it is the most widely used strain in mechanistic studies. In this model, paresis begins in the hindlimbs and spreads rostrally to the forelimbs. Males experience earlier onset, greater disease severity, and shorter survival than females. However, the influence of sex on patterns of declining motor function between forelimbs and hindlimbs as well as among distinct, spinal-innervated muscle groups within each limb are not fully understood. To provide a higher resolution framework of degenerating motor function across the body, we conducted more comprehensive, limb-dependent and independent measures of motor decline over the course of disease. Subsequently, we compared the timing and intensity of these features across sex, and we consider to what extent these patterns are conserved in clinical observations from human ALS patients. We found male mice experienced earlier and less localized onset than females. We also report distinct motor features decline at different rates between sexes. Finally, mice showed differences in correlation between the decline of left- and right-side measures of the hindlimb. Consequently, our findings reinforce and refine the utility of the SOD1 mouse in modeling more highly resolved, sex-specific differences in ALS patient motor behavior. This may better guide preclinical studies in stratifying patients by sex and anatomical site of onset.

## Introduction

Amyotrophic lateral sclerosis (ALS) is a progressive disease in which upper and lower motor neurons rapidly degenerate. In humans, its onset is 49.7+/-12.3 years in North America, starting with focal weakness, progressing to involve most muscles, and usually causing fatal respiratory failure within 5 years (1).

Approximately two-thirds of ALS patients experience spinal onset, developing focal muscle weakness or wasting in one or more limbs first; the remaining majority experience bulbar onset, first exhibiting difficulty speaking, breathing, or swallowing (1). Before the age of 65, men are at higher baseline risk than women for developing ALS and are likelier to experience spinal onset of motor dysfunction, including cases initially beginning in the upper or lower extremities, while women are likelier to experience onset in bulbar regions (2–5). However, the influence of sex on the patterns of declining motor function between arms and legs as well as among distinct, spinal-innervated muscle groups within each limb are not fully understood (6–8). Approximately 90% of ALS cases are sporadic (sALS), without a family history or pathogenic genetic variants; the other cases are familial (fALS) (9). The most common identified genetic cause is the *C9orf72* expansion (∼40% of fALS cases); the second most common is *SOD1* mutations (∼20% of fALS cases) (10). After *SOD1* became the first causative gene discovered for ALS, the first animal model developed was the SOD1G93A ALS mouse model (hereafter referred to simply as SOD1 mice) (11,12), and it remains the most widely used model for preclinical studies.

Despite ostensible differences between quadrupedal rodent and bipedal human gait patterns, the conceptual basis for gait analysis remains analogous across species (13), and the effects of ALS in this mouse model on biomechanical components highly resemble the effects of ALS in patients (14). For example, males experience onset earlier than females, and sex does not affect disease duration (15–17). However, there are many aspects of this model which differ from ALS patients. Among these, SOD1 mice, specifically in the C57BL/6 background, have a stereotypical onset location, starting with declining hindlimb motor function, subsequently progressing towards anterior body parts (18). While this enables a tractable model for mechanistic and therapeutic research, it contrasts with the variable onset locations of motor dysfunction in patients.

Furthermore, prior studies using SOD1 mice have extensively characterized sex differences in hindlimb motor function decline. However, fewer studies have considered forelimb decline in relation to hindlimb decline and have not directly compared their motor features on the same measurement scale (17,19,20) (21–23).

Here, we established a multimodal (bodyweight, strength, function) comparison of ALS-induced forelimb, hindlimb, and limb-independent motor dysfunction progression between SOD1 and wild type (WT) mice using several behavior tests and evaluated the effect of sex on functional degradation. We found males experienced earlier and less localized onset than females. Unlike patients, some mouse motor aspects decline at significantly different rates between sexes, as well as between different sides of the body (24,25).

Consequently, these findings reinforce and refine the SOD1 mouse’s utility in modeling more highly resolved, sex-specific differences in ALS patient motor behavior. These parallel and contrasting phenotypes based on sex and onset location may help guide pre-clinical studies aimed at developing treatments for ALS patients.

## Results

### Male SOD1 ALS mice experience greater and earlier decline in body weight and limb function

We quantified body weights for male and female SOD1 ALS and WT C57BL/6 mice at several time points between P25 and P150+ (Fig. 1A and S1A). The body weight of SOD1 mice averaged lower than WT mice of the same sex by P90 (Fig. 1B). When evaluating SOD1 population body weight average as a percentage of WT body weight average from the same sex and time point, we considered a five percent reduction as meaningful (26). Average male SOD1 body weight change expressed this way exhibits greater than five percent reduction by P90, with further reduction in subsequent timepoints (Fig. S1B). Of note, average female SOD1 body weight exhibited greater than five percent reduction at P25; however, it recovered to two percent reduction by P50, then progressed to greater than five percent reduction in subsequent timepoints starting at P90. This may indicate a sex-based effect on postnatal development rather than muscle atrophy in early adulthood. Overall, these data indicate that male transgenic mice experience a more severe body weight decline over the course of the disease than females.

**Fig. 1.**
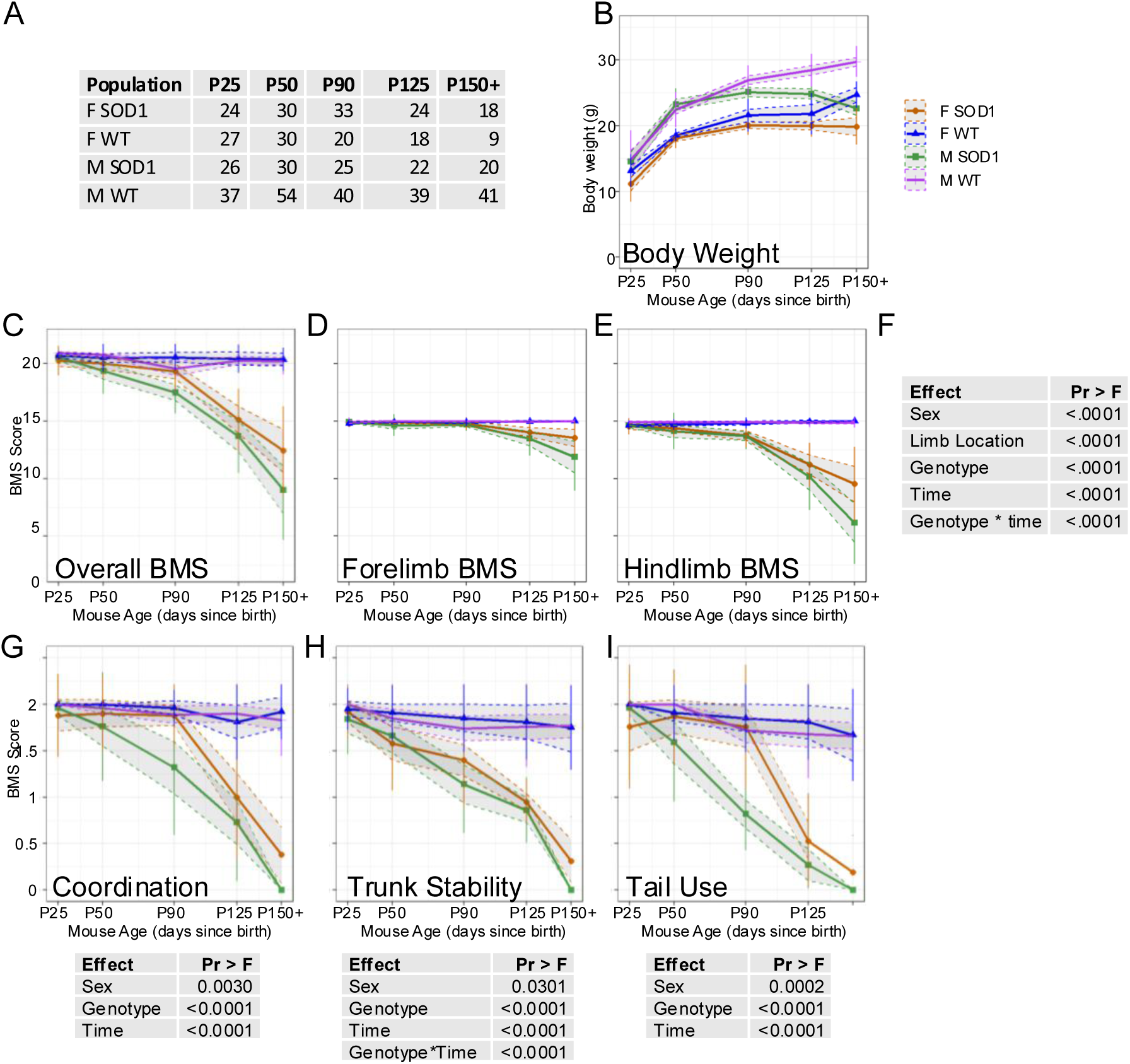
Male SOD1 mice experience a greater and earlier decline in limb and non-limb-specific functions. Average values for each sex-genotype population over five timepoints. A) Number of mice represented at each data point. B) Body weight progression. C-E) Overall, forelimb-only, and hindlimb-only BMS score progression. F) Linear mixed model of BMS scores with repeated measure analysis to determine p-values. G-I) Non-limb-specific BMS metrics: coordination, trunk stability, and tail use, as well as generalized linear mixed effects models with random intercept to determine p-values for each metric.

We also adapted a measure of locomotor recovery after spinal cord injury, the Basso Mouse Scale (BMS) (27) for our observational time course for ALS-induced motor decline (Table 1). We found SOD1 BMS scores averaged significantly lower than WT by P50, and then decayed significantly in successive timepoints (Fig. 1C). When evaluating SOD1 population BMS average as a percentage of WT BMS average from the same sex and time point, a reduction of greater than five percent is observed at P50 for males and P90 for females (Fig. S1C). Separately considering forelimb and hindlimb BMS, male SOD1 mice experienced more severe decline than females in both regions (Fig. 1D and 1E). Furthermore, average SOD1 hindlimb BMS scores degraded faster than forelimb scores in both sexes by P125, whereas WT mouse BMS rankings did not change significantly (Fig 1D and 1E). When evaluating SOD1 population forelimb BMS average as a percentage of WT forelimb BMS average from the same sex and time point, a reduction of greater than five percent is observed at P125 for both sexes (Fig. S1D). However, a reduction of greater than five percent is first observed in hindlimbs at P50 for male SOD1 mice, whereas female SOD1 mice exhibit this decrease at P90 (Fig. S1E). A linear mixed model with repeated measures generated with these data indicated several effects on overall BMS score decline (Fig. 1F). Among them, sex was a prominent feature: male SOD1 mice experienced more severe decline of movement, consistent with the body weight phenotype. These sex differences are not evident when quantifying the rate of decline for overall, forelimb, and hindlimbs using a simple linear slope (Figs. S2A and S2B). This observation underscores a non-linear pattern of motor decline in SOD1 male and female mice. Overall, these data show that hindlimb function declines faster than forelimb function in the SOD1 mouse model, and male mice experience more severe decline than females in these aspects.

**Table 1.**
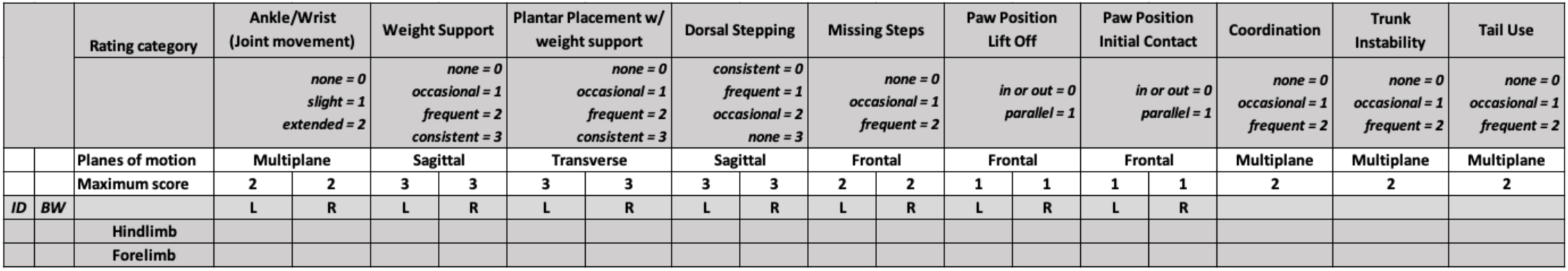
Scoring criteria of the modified Basso Mouse Scale. Joint movement: ankle/wrist exhibits full (2), less than half (1), or no (0) range of motion during stepping. Weight support: knee/elbow joints consistently (3), frequently (2), occasionally (1), or never (0) extend and avoid touching ground; hindquarters/forequarters are fully (3), half (2), quarter (1), or not (0) raised off the surface. Plantar placement with weight support: paw’s digits are in complete extended position and first and last digits contact the ground so its plantar surface consistently (3), frequently (2), occasionally (1), or never (0) appears flat on the ground during the stance phase. If no weight-supported stepping or if toe drag occurs, 0. Dorsal stepping: paw’s dorsal surface never (3), occasionally (2), frequently (1), or consistently (0) occurs with weight support. Missing steps: no (2), occasional (1), or frequent (0) lapses in forelimb-hindlimb coordination during abbreviated walking bouts, or failure to maintain weight support during stance. Paw position: paw’s middle digits are parallel (1) or rotated (0) in comparison to the body’s long axis. Scored twice, once at lift off, once at initial contact. Coordination: ≥3 (2), <3 (1), or no (0) assessable passes performed at constant speed. Per each forelimb step, a hindlimb step is never (2), occasionally (1), or frequently (0) not taken. Contralateral limbs alternate consistently (2), alternate occasionally (1), or frequently don’t alternate (0) per gait cycle. Trunk instability: 2 points- trunk doesn’t lean/sway, tail’s distal third is steady and elevated, and there are no postural deficits; 1- pelvis/haunches predominantly dip, rock, or tilt, there is some sway in the hindquarters and/or the tail’s distal third is not steady; 0 - severe postural deficit e.g. pronounced lean, waddling, near- collapses of the forequarters/hindquarters, hunch, events of the hindquarters contacting the ground, neck tilt, dropped head and/or torso; signs of one or more immobilized joints, curled limb(s), or inverted hind limbs (facing upward). Tail position: tail’s entire length remains elevated during walking (2); tail is held up least once during locomotion or 2/3 of the tail length is consistently limp (1); tail’s base is consistently dragged (0).

### Coordination, trunk instability, and tail use decline more severely among SOD1 males

In addition to limb-specific BMS measures, we also compared differences in non-limb-specific BMS measures. Coordination is defined as: for every forelimb step, a hindlimb step is taken and the hindlimbs must alternate (27). This measure began degrading in SOD1 males by P50 and in SOD1 females by P125 (Fig. 1G and S1F).

Trunk instability began degrading in SOD1 males by P25 and in SOD1 females by P50 (Fig. 1H and S1G). Tail use began degrading in SOD1 males by P50 and in SOD1 females by P125 (Fig. 1I and S1H). In considering SOD1 population BMS score average as a percentage of WT score average of the same sex and time point, the consistent decline of coordination, trunk instability, and tail use starts at an earlier timepoint for SOD1 males than for SOD1 females (Fig. S1F-H). However, once average SOD1 female tail use began declining, it decreased faster than it did in SOD1 males (Fig. S1H). Incidentally, we observed multiple SOD1 mice with tail use scores of 0-1 lift their tail bases while inside their cage enrichment tube rotated more than ten degrees, suggesting that lack of tail suspension in pathogenic mice owes to preference, not incapability. Interestingly, SOD1 males exhibited significantly steeper rate of decline than SOD1 females in all three metrics when considering a linear slope (Fig. S2C), again underscoring the non-linear differences in patterns of motor decline between sexes. Sex, genotype, and timepoint significantly affected limb-independent motor scores when analyzed using linear mixed models with repeated measures (Fig. 1G-I). Similar to body weight, coordination and tail use scores in SOD1 females averaged significantly lower than WT scores at P25, though this relationship recovers to less than a five percent difference by P50 and subsequently declines rapidly by P125 (Fig. S1F, and S1H); this may indicate a sex-based, postnatal motor function difference preceding later motor function decline. Notably, trunk instability exhibits the earliest decline in SOD1 mice for both sexes; males at P25 and females at P50. This decline occurs before the greater than five percent decrease in body weight observed at P90. All other features considered thus far (forelimb and hindlimb BMS, coordination, and tail use) exhibit decline at or after P90 in female SOD1 mice. In contrast, all other features considered thus far, except forelimb BMS, exhibit decline before P90 in male SOD1 mice. Altogether, these data reveal limb- and non-limb-specific measures of spinal motor function decline prior to P90, the age generally regarded as ALS onset in SOD1 mice, and these functions decline more severely in males.

### Limb-specific motor functions decline faster in SOD1 males

Since BMS is a composite score of multiple movement features, we compared each feature within limb-specific measures to resolve their respective contributions to the observed sex differences (Fig. 2). In each limb-specific BMS criterion, average onset of decline in SOD1 males occurred either earlier or at the same time point as females. Average onset occurred earlier in SOD1 males for eight criteria (forelimb: missing steps, paw position, hindlimb: weight support, plantar placement at P50, and forelimb and hindlimb: joint movement, dorsal stepping at P150). Among SOD1 females, forelimb joint movement and hindlimb dorsal stepping never declined. By P150, each limb-specific BMS metric averaged worse among SOD1 males than SOD1 females.

**Fig. 2.**
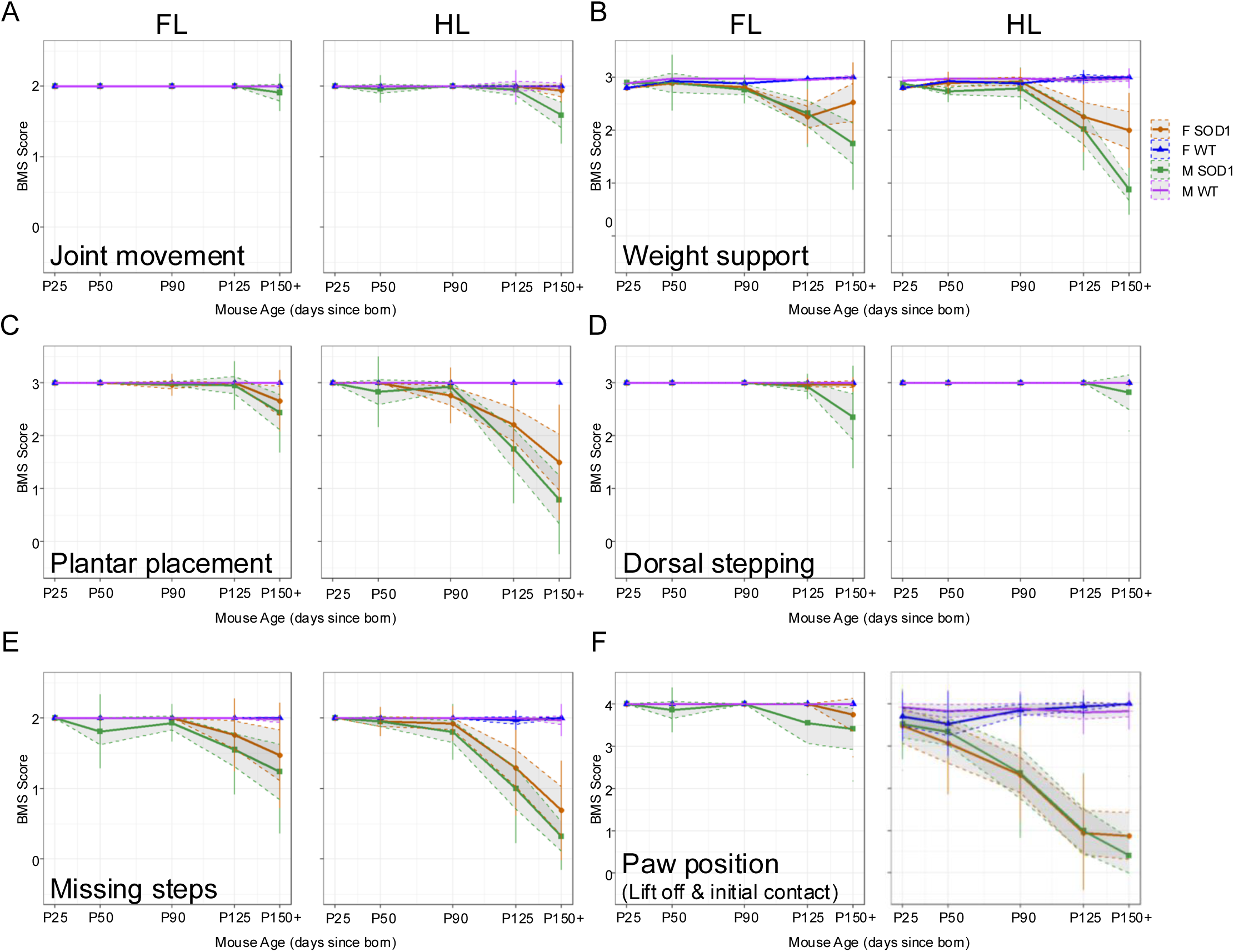
Limb-specific BMS metrics decline more severely in SOD1 males. Limb-specific BMS metrics in forelimb (FL) and hindlimb HL. A) Joint movement. B) Weight support. C) Paw placement. D) Dorsal stepping. E) Missing steps. F) Paw position at lift off and initial contact.

Notably, a decrease in average hindlimb paw position in both male and females is observed at P25 and continually decreases until endpoint, indicating it as the first BMS measure to be affected. From these measures, we conclude that hindlimb weight support, plantar placement, and paw position are the movement features that contribute the most to the reduced hindlimb BMS scores by P50.

### Grip strength declines earlier in SOD1 males but faster in females

To assess another measure of forelimb function, we collected grip strength at the same time points as body weight and BMS between P25 and P150+ (Fig. 3A). Normalized to body weight, average SOD1 grip strength monotonically decreased at every subsequent time point for both sexes (Fig. 3B). A generalized linear mixed model with repeated measures applied to these data indicated several significant effects (sex, genotype, time, and time-genotype interaction) on grip strength decline (Fig. 3C). At P25, SOD1 and WT males had similar grip strength, but by P50, WT males increased in grip strength while SOD1 males did not (Fig. 3B), which reflected a greater than five percent reduction at P50 (Fig. S1I). Grip strength continuously declined throughout adolescent, adult, and end stages in the SOD1 male mice (Fig. 3B). These data are consistent with the earlier onset of forelimb missing steps and paw position deficits in SOD1 males at P50 (Figs. 2E and 2F). At P25, SOD1 and WT females had similar grip strength, and both genotypes showed greater normalized grip strength than males (Fig. 3B). However, SOD1 females showed a faster overall rate of grip strength decline than SOD1 males (Fig. 3D), despite reaching a more delayed threshold of five percent reduction from WT females at P90 (Fig. S1I). Altogether, these data reveal different kinetics of grip strength between sexes and genotype throughout the course of disease progression.

**Fig. 3.**
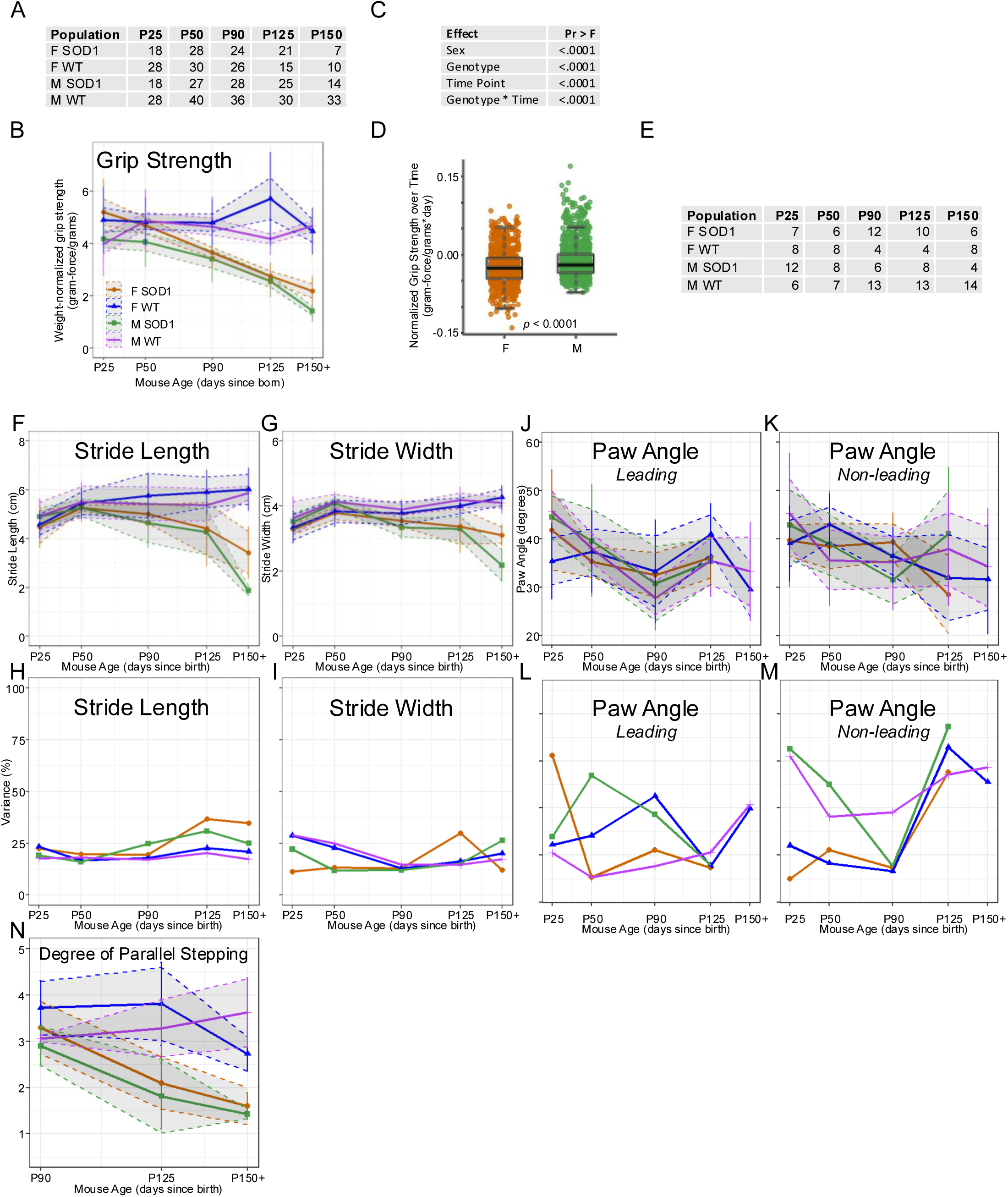
Female SOD1 mice experience a faster loss of grip strength, but male SOD1 mice exhibit the worst grip strength and footprint measures by end stage. A) Number of grip strength-tested mice at each timepoint. B) Grip strength force progression. C) Generalized linear mixed effects model with random intercept used to determine p-values. D) Rate of change in grip strength per day in ALS populations. A two-tailed t-test between males and females was applied. E) Number of ink pawprint-tested mice at each timepoint. F) Stride length progression. G) Stride width progression. H) Stride length variance. I) Stride width variance. J) Leading paw angle progression. K) Non-leading paw angle progression. L) Leading paw angle variance. M) Non- leading paw angle variance. N) Degree of parallel stepping progression, determined by dividing the wider angle of both paws in the same gait cycle by the narrower angle. Each subject has two paw angle data points at each timepoint, the average angle of the paw leading gait and that of the paw not leading gait.

### Male SOD1 mice exhibit worse footprint test measures by end stage

To assess other measures of hindlimb function, we collected ink paw print data at the same time points between P25 and P150+ (Fig. 3E). Average WT stride length and width increased in nearly every successive time point in both sexes, and WT measures averaged higher than their SOD1 counterparts by P90 (Fig. 3F and 3G). Average SOD1 stride length and width continually decreased after P50, and after P125, these measures in SOD1 males were lower than SOD1 females (Figs. 3F and 3G). Whereas WT mice had low variance in their stride lengths throughout the observed ages, SOD1 mice showed slightly higher variance in their stride lengths beginning at P90 (Fig. 3H). Conversely, WT mice showed slightly higher variance than SOD1 mice in their stride width prior to P90; SOD1 females showed an acute increase in stride width variance at P125 then reduced to below WT at P150+ (Fig. 3I), likely due to reduced sampling at these time points (Fig. 3A). Relative to females, SOD1 males showed a delayed increase in stride width variance by P150+ (Fig. 3I). Leading and non-leading paw angle measurements for all populations were highly variable at all time points (Figs. 3J-M).

Finally, we analyzed the degree of parallel stepping. A reduced ratio between left and right signifies the placement of hindlimbs in a hopping-like pattern, and SOD1 mice of both sexes exhibit this pattern at P125 and end stage (Fig. 3N). Consistent with reduced coordination in BMS data, these data suggest deficiencies in the central pattern generation. Altogether, ink paw print data concurred with the BMS data, demonstrating worse hindlimb functional decline in gait and step cycle patterning in SOD1 mice with less overt differences between male and female mice.

### Motor decline rates across contralateral limbs are less correlated in male SOD1 mice

When aggregating forelimb and hindlimb scores, there was no significant difference between left and right limb BMS score decline for any sex-genotype population (Figs. 4A and 4B). However, when left and right limb scores were separated between forelimb and hindlimb, the left- and right-limb correlations of some declining BMS measures differed significantly between sexes (Figs. 4C-4E). Whereas no significant differences were observed in forelimb between sexes (Fig. 4C), decline rates in left and right hindlimb weight support and hindlimb missing steps were less correlated among SOD1 males (Fig. 4D). This suggests that pathogenic factors responsible for these aspects of motor decline spread slower between hindlimbs in SOD1 males.

**Fig. 4.**
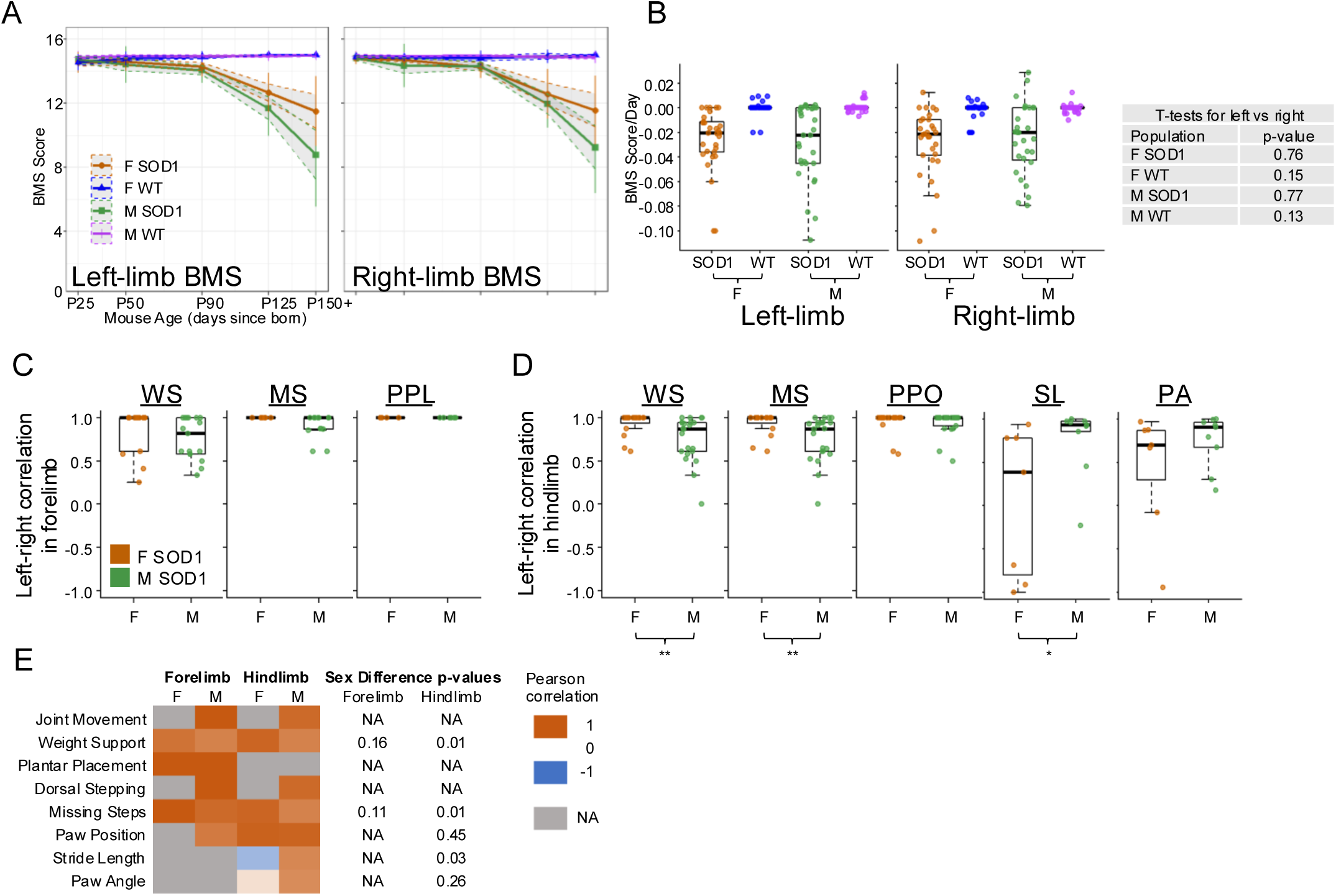
There is a sex difference in the relative weight support decline of left and right hindlimbs. A) Left- and right-limb-separated BMS score progression. B) Rates of left- and right-limb score BMS change for subjects with at least two measurements. C-D) Box plots of correlation between left- and right-forelimb and hindlimb rates of decline for limb-specific BMS scores and ink pawprint tests. C) Forelimb weight support (WS), missing steps (MS), and paw placement (PPL). D) Hindlimb weight support (WS), missing steps (MS), paw position (PPO), stride length (SL), and paw angle (PA). E) Correlations between left- and right-forelimbs and hindlimbs for each sex. Each panel represents the average Pearson’s correlation of the values shown in C and D. “NA” comparisons occurred when no subject in one of the populations exhibited change over time in either left- or right-limb score, which occurred for each criterion. P-values of sex differences in left-right correlations were determined via Wilcoxon rank-sum test.

Conversely, decline rates in left and right hindlimb stride length were less correlated among SOD1 females. These features are consistent with reduced coordination in both sexes, which may contribute to this observation. Taken together, these analyses highlight distinct sex patterns of how declining motor function correlate across the midline over the course of ALS in SOD1 mice.

### Coordinated decline of motor function measures reveals four distinct sex-conserved clusters

To assess how each of the motor behavioral measures relate to each other, we correlated each metric with each other and compared the distribution of correlations between sexes (Fig. S3). In this manner of observation, one correlation that significantly differed was the correlation between declining forelimb missing steps with grip strength. This correlation was significantly higher in SOD1 females. This suggests that motor units involved with weight support during initial contact and stance phases are more interlinked with those involved with grip strength in females. To observe higher order patterns of decline among all the motor features we measured, we performed hierarchical clustering of SOD1 ALS-induced decline in each BMS criterion, forelimb grip strength, stride length, and stride width (expressed as the percent of WT scores of the same sex and time point). This analysis revealed distinct degradation patterns between sexes (Fig. 5). Among females, measures in cluster 1 (trunk stability, tail base height, coordination, hindlimb paw position) experienced little degradation in early timepoints, but more severe degradation than other measures by P125-150. Cluster 2 measures (weight support, missing steps, hindlimb stride width) experienced nearly no degradation in earlier timepoints, but some degradation in P125-150. Cluster 3 measures (dorsal stepping, joint movement, forelimb plantar placement, forelimb paw position) experienced little degradation. Cluster 4 measures (hindlimb stride length, hindlimb missing steps, hindlimb plantar placement, forelimb grip strength) started degrading by P90 and worsened more severely than cluster 2 towards P150, though not as severely as cluster 1 measures.

**Fig. 5.**
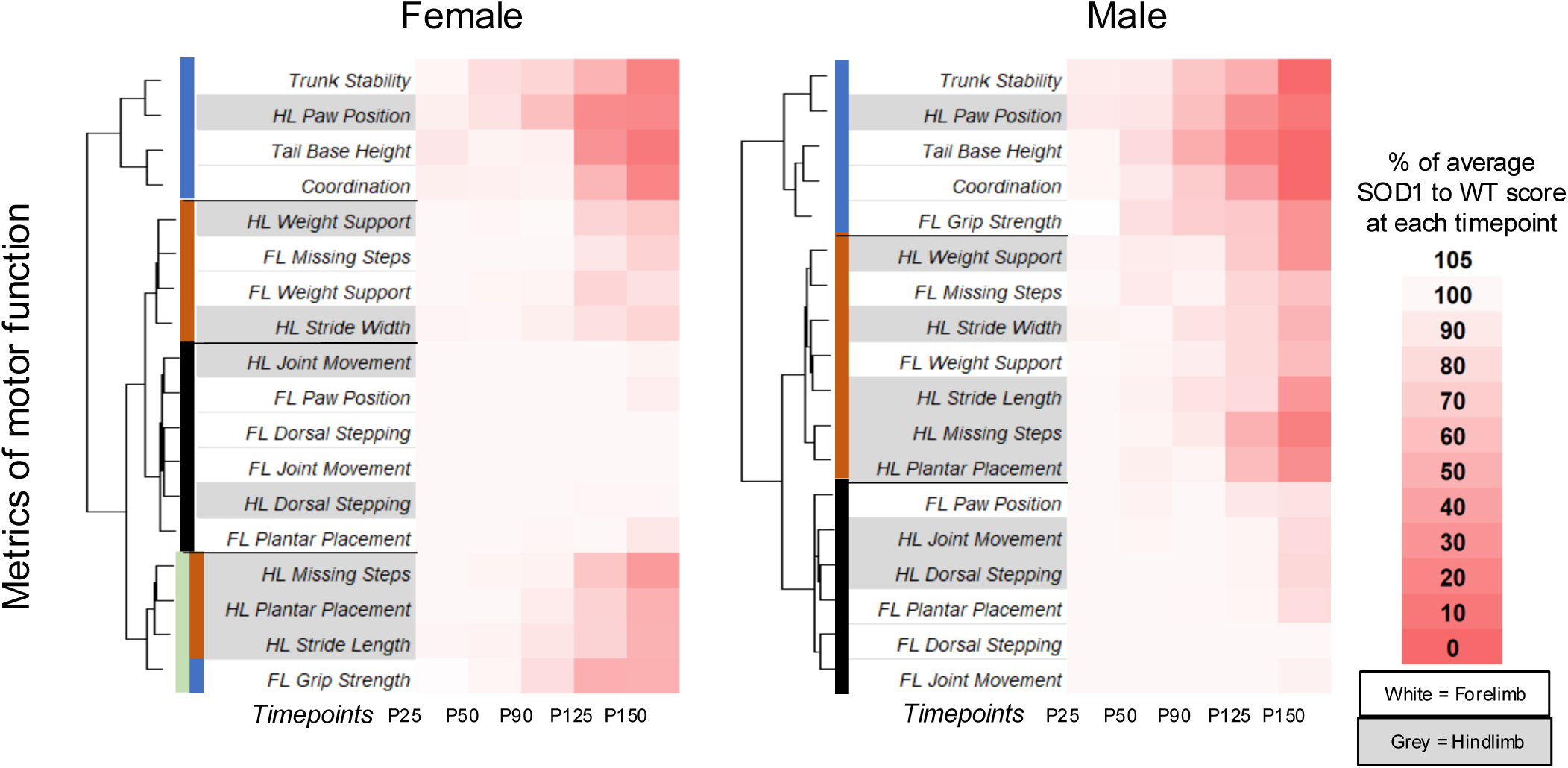
Motor functions decline in sex-specific patterns. Comparison of SOD1 mouse sex-specific patterns of the changes over time in individual BMS metrics, grip strength, and ink pawprint measures, expressed as percentages of WT scores of the same sex and timepoint. Dendrograms were constructed using Ward’s hierarchical clustering method, with Euclidean distance as the metric for measuring similarity between data points. In males, there are three major clusters of metrics that decline in SOD1 mice with similar patterns, and these are labeled with three colors: blue, red, and black. These three clusters of metrics are largely conserved in female SOD1 mice, however some metrics from the blue and red clusters in males form one cluster in the females: light green. All clusters of motor functions in males and females exhibit distinct patterns of decline considering age of onset and percentage decrease from WT mice.

Among males, cluster 1 is well-conserved, containing all the measures as females but also included forelimb grip strength. However, this group of measures experienced earlier onset of degradation than in females.

Clusters 2 and 4 (without forelimb grip strength) formed a single “supercluster” in males, describing an earlier and more gradual decline (as opposed to a sharper drop at P90) than observed for females. Cluster 3 was fully conserved in males; however, these measures exhibited more severe decline by end stage. Overall, the distinct clustering patterns among declining motor functions in SOD1 mice suggests motor groups may have differential interlinked dependencies or pathways of spreading pathology between sexes. Males exhibited earlier onset, more interlinked motor units, and greater severity of declining motor function.

## Discussion

The transgenic SOD1G93A mouse model stereotypically and reproducibly exhibits the pathology of ALS; therefore, it is valued as the workhorse model for ALS pathogenesis in basic and pre-clinical studies. To better track the spread of degenerating motor functions across the body axis, we conducted more comprehensive, direct comparisons of forelimb and hindlimb ALS-induced motor decline. As the integrated pattern of walking in ALS goes awry, the biomechanical complexes of the three body planes (*transverse, sagittal, frontal)* change dynamically; these behaviors include rhythmics of gait, interlimb coordination, spatial and timed symmetry of walking pattern, stretch reflexes, balance, and posture maintenance (28). While several objective and qualitative techniques such as 3D motion capture systems and imaging have been developed and applied to biomechanical studies in animal models, the BMS scale remains reliable, because it describes multiplane motion in a freely behaving environment (29).

Our findings from these assays reinforce the understanding that SOD1 ALS-induced motor neuron dysfunction occurs earlier and progresses faster in male than female mice. One exception to this trend is motor performance measured through the grip strength assay, where males experience earlier onset of decline, but females show faster overall rate of decline. Clinically, younger ALS patients tend to be male, and onset of fine motor impairment in spinal-onset patients occurs earlier in males (3,5). Contrastingly, the rate of motor decline, assessed by the revised ALS functional rating scale (ALSFRS-R) has been reported to be faster in female patients, regardless of onset site (30). However, ALSFRS-R is a composite score of multiple measures of gross motor performance, and this aggregate comparison may potentially mask contradictory differences among individual ALSFRS-R metrics. Therefore, stratified comparisons for these individual features, analogous to the individual BMS features we have analyzed, may reveal sex differences more accurately.

Additionally, our analyses revealed that the onset of dysfunction in motor groups are less localized, or more widespread in males. This observation motivates deeper investigation into focused motor group data in ALS patients to see if similar sex differences are conserved. Our BMS data also showed postural control, assessed by trunk instability, experiences the earliest impairment among all other BMS measures in both males and females. In patients, trunk instability has been proposed as a sensitive indicator of ALS severity, closely correlated with motor function and balance test scores (31). This observation in mice may yet warrant closer attention to reduced postural motor control as a sign of onset.

Across both sexes, hindlimb paw position was the earliest element of the limb-dependent BMS score to decrease. We noted that external rotation of the hindpaw was the most misaligned angle relative to the body’s long axis, which was evident shortly after the propulsive period ended and the swing phase was initiated. This phenotype results from failed foot inversion, signifying dysfunction of the tibialis posterior, flexor digitorum longus, and flexor hallucis longus via the tibial nerve. Decrease of BMS plantar placement of the hindlimb, which results from failed dorsiflexion, shortly followed failing hindlimb paw position. This suggests that the dysfunction of the tibialis anterior flexor digitorum longus, and flexor hallucis longus via the deep fibular nerve follows dysfunction of the tibial nerve. In lower limb onset ALS patients, ankle dorsiflexion weakness, signified by foot drop, is a common clinical manifestation during the early stages of disease (32,33). While weak foot inversion, plantarflexion, and hip abduction can also contribute to altered gait mechanics and foot drop-like presentations, these dysfunctions are typically secondary features and become more apparent as ALS progresses (32). Therefore, muscles responsible for ankle dorsiflexion are regarded as the earliest groups to decline in most of these ALS patients. A possible explanation for this difference between mice and humans may be that the muscles innervated by the tibialis posterior experience higher usage and stress due to the midfoot strike pattern in mice, opposed to the heel strike in humans. Indeed, the lateral cuneiform is fused to the navicular to provide more stability (34), limiting abduction of the forefoot at the midfoot, and thus cause the entire foot to rotate outward because of a weakened tibialis posterior. In light of our observations in SOD1 mice, the onset of tibial nerve deficits in these patients may nevertheless merit closer investigation.

Markedly, analysis of limb-dependent features in SOD1 mice showed differences in correlation between the decline of left- and right-side measures of the hindlimb. Males exhibited lower correlation of weight support, which depends on the activation of antigravity muscles such as the quadriceps and gastrocnemius, and missing steps, which signifies dysfunction of muscles responsible for initiating and coordinating steps. Females exhibited lower correlation of declining stride length, signifying dysfunction of several hindlimb muscles. Prior analyses of hand-held dynamometry, Tufts Quantitative Neuromuscular Exam, and Accurate Test of Limb Isometric Strength protocols in ALS patients reported high correlations of progression in declining muscles across left and right lower limbs (24,25). However, neither study accounted for sex. Considering our findings, these patient data warrant further investigation for sex-effects.

Our interpretations and conclusions may be limited by several factors. While our analysis of five time points revealed multi-factorial effects on motor dysfunction, additional time points could improve our resolution of ALS progression in each feature. Consistent sampling per mouse and time point could also provide more robust longitudinal analyses. Furthermore, we investigated the high-copy number SOD1G93A model in the C57BL/6J strain, whereas other SOD1 transgenic models in multiple other background strains have reported variations in disease phenotypes. For example, the low-copy number SOD1G93A model exhibits an earlier onset of reduced grip strength in females, but a faster rate of progression in males (35). Overall, SOD1 mutations account for 20% of familial ALS cases and 2% of all ALS cases. Expanding the application of our motor assays to other genetic mouse ALS models can reveal common patterns of onset and declining motor function to offer better physiological underpinnings of ALS, and perhaps enable the development of more targeted or universal therapies.

## Acknowledgements

We would like to acknowledge the Cedars-Sinai Biobehavioral Core and thank Dr. Catherine Bresee for helpful advice on statistical analyses. This study was supported by grants from the National Institute on Aging (R00AG056678), Center for Research in Women’s Health Sciences Research Award by the Louis B. Mayer Foundation, and Winnick Foundation Research Scholars Award.

## Author Contributions

O.S. designed and performed experiments and data collection. I.T. performed data collection and analysis and figure generation. M.L. performed data analysis. A.D. performed data collection and analysis. R.H. directed the research, designed experiments, and provided funding. O.S., I.T., and R.H. wrote the manuscript, and all authors provided edits prior to submission.

## Data Availability

All source data are provided with this paper in Supplemental Table 1.

## Code Availability

The code utilized for our analysis are available at https://github.com/ritchieho.

## Methods

All procedures were approved by Cedars-Sinai Medical Center’s Institutional Animal Care and Use Committee (protocol 7496). We compared motor function of transgenic mice (N = 58) from a single strain with a SOD1 ALS mutation with WT mice (N = 64) at 5 time points: 25, 50, 90, 125, and 150+ days after birth. This strain of mice was bred from a B6.Cg-Tg(SOD1*G93A)1Gur/J with C57BL/6 background. Initially, C57BL/6J female mice were obtained from the Jackson Laboratory. The animals were bred in harems using optimal outbreeding strategy to produce experimental animals. Transgenic and corresponding non-transgenic littermates were used to create five timepoint groups per sex to perform behavior analysis. C57BL/6 were used as controls.

Genotyping for the presence of mutant SOD1 gene as well as sex was contracted through Transnetyx using their standard probe sets for the SOD1 gene and Y chromosome. No sample size calculations were performed for any assay. Mice that failed neurological scores were excluded. Assessors were aware of subjects’ sex, age, and genotype during each assay.

We evaluated body weight, as it is a strong prognostic factor for ALS onset (16,36,37). We applied three mouse model functional analyses to evaluate the motor neuron function of mice: modified Basso Mouse Scale, hand grip strength evaluation, and footprint analysis. Each mouse was evaluated at 25, 50, 90, and 125 days of age, and at its endpoint occurring at or after 150 days. SOD1 mice were monitored until humane endpoint, assessed by loss of righting reflex: inability of the animal to right itself to maintain normal posture when manually placed on its side (dorsal side of the scapular and lateral side of the hip contact the floor) within 30 seconds, or paralysis causes other concerns for the animal’s general welfare or its capacity to eat/drink and ambulate. This assessment is performed free of obstacles, on bedding in the animal’s cage environment.

### Basso Mouse Scale

The Basso Mouse Scale for Locomotion (BMS) is used to evaluate motor neuron function of experimental mice following spinal cord injuries, conducted by visual observation of a subject walking on an open tray for three minutes and scoring its motor function along several distinct criteria (27). The BMS measures multiplanar (coronal, sagittal, and transverse) motion. It is a binary scale (presence [1] vs absence [0]) based on visual assessment of seven locomotor attributes: ankle movement, frequency of dorsal and plantar stepping, weight support maintenance, coordination of gait, paw position at initial contact and lift off, trunk instability, and tail function control. The original BMS score range is 0-9 with subscores; nine points indicates absence of motor neuron dysfunction. Originally, the BMS was used to rate motor function recovery of hindlimbs affected by spinal cord injury (SCI) in mouse models. For our investigation, the BMS evaluated progression of motor neuron dysfunction in SOD1 transgenic mice compared to WT mice. We converted the original BMS scale into a ratio scale of 21 scores to evaluate twelve behavioral categories of the muscle groups function during disease progression. During each evaluation we rated each locomotion criterion with independent scores per limb, and we established each subject’s total BMS score via averaging their limbs’ scores.

The original BMS’s criterion ordering is ranked by degree of severity, but intervals between the scale points may be inconsistent (27). We therefore converted the original BMS to a ratio scale to better evaluate gradual dysfunctional development in muscle groups during ALS progression; the 0 point and numerical relationships between values are meaningful throughout the scale. This allowed us to define suitable statistical tests to describe sex and anatomical location differences at different time points (38). The highest and lowest point scores for each criterion indicate total presence and absence of its locomotor attribute during gait, respectively. An ‘assessable pass’ occurs whenever the subject travels continuously and straight for a distance of at least three body lengths. ‘Occasional’ is defined as occurring less than half the time; ‘frequently’ as more than half; and ‘consistent’ as over 95% of the time. The modified scale criteria are defined thusly:

a. Joint movement in the limb. 2 points- ankle/wrist joint exhibits full range of motion; 1- joint exhibits less than half its full range of motion during stepping; 0- joint exhibits no motion.
b. Weight support (ability to elevate hindquarters/forequarters throughout the stance). 3 points- knee/elbow joints continuously extend and do not touch ground, and hindquarters/ forequarters do not contact the surface at all; 2- joint frequently does not touch ground, hindquarters/forequarters are half raised off the surface; 1- joint frequently touches ground, hindquarters/forequarters are 1/4 raised off the surface; 0- joint consistently touches ground, hindquarters/forequarters are not elevated.
c. Plantar placement with weight support. 3 points- paw’s digits are in complete extended position and first and last digits contact the ground so its plantar surface consistently, 2- frequently, 1- occasionally, or 0- never appears flat on the ground during the stance phase. If no weight-supported stepping or if toe drag occurs, 0.
d. Dorsal stepping. 3 points- paw’s plantar surface provides weight support to every step in assessable passes; 2- paw’s dorsal surface occasionally provides weight support during assessable passes; 1- paw’s dorsal surface frequently provides weight support during assessable passes; 0- paw’s dorsal surface consistently provides weight support during assessable passes, or no weight-supported stepping occurs.
e. Missing steps (dorsal or plantar), defined as frequency of re-established weight support stepping. 2 points- no missing steps (constituting lapses in forelimb-hindlimb coordination during abbreviated walking bouts, or failure to maintain weight support during stance) occur; 1- occasional missing steps; 0- frequent missing steps.
f. Paw position. At lift off: 1 point- paw’s middle digits are parallel to the body’s long axis; 0- paw’s middle digits are rotated, pointing internally (inward) or externally. At initial contact: 1 point- paw’s middle digits are parallel to the body’s long axis; 0- paw’s middle digits are rotated.
g. Coordination. 2 points- mouse can perform three assessable passes at constant speed and a distance of at least 3 body lengths, a hindlimb step is taken for every forelimb step, and contralateral limbs alternate in each assessable pass; 1- less than three assessable passes at constant speed occur, hindlimb steps occasionally aren’t taken per forelimb step, and/or homologous limbs don’t alternate in each gait cycle; 0- no assessable passes occur at constant speed and/or hindlimb steps frequently aren’t taken per forelimb step, and there is a loss of alternating gait.
h. Trunk instability (ability to maintain postural control and stable trunk during walking). 2 points- trunk doesn’t lean/sway, tail’s distal third is steady and elevated, and there are no postural deficits; 1- pelvis/haunches predominantly dip, rock, or tilt, there is some sway in the hindquarters and/or the tail’s distal third is not steady; 0 - severe postural deficit e.g. pronounced lean, waddling, near-collapses of the forequarters/hindquarters, hunch, events of the hindquarters contacting the ground, neck tilt, dropped head and/or torso; signs of one or more immobilized joints, curled limb(s), or inverted hind limbs (facing upward).
i. Tail position. 2 points- tail’s entire length remains elevated during walking; 1- tail is held up at least once during locomotion or 2/3 of the tail length is consistently limp; 0- tail’s base is consistently dragged.

Subjects were tested on an open field for 3 minutes while trotting (taking a hindlimb step with each forelimb step) or walking (three feet simultaneously in stance, with a distinct alternating inter-limb coordinated order) (39). Subject gait is approximately 90% trotting; subjects generally walk abnormally when trotting becomes unfeasible due to impaired coordination.

### Grip strength evaluation

Hand grip strength is a known indicator of age-related decline in mice (40), and it has been used to evaluate the SOD1G93A mouse model, as it reflects mouse forelimb muscle condition (41). At each timepoint, subjects were manually lowered onto the top bar of a grip strength meter grid (San Diego Instruments, San Diego, CA, USA) angled at 40 degrees. Once grip is established, the subject is pulled back slowly and steadily until it releases its grip. Evaluations obtained gram-force measurements of peak grip strength, repeated three times with 20 seconds between each trial for each forelimb independently (the other prevented from obtaining grip by wrapping tape around its wrist) and for both forelimbs together.

### Footprint test

Stride length and stride width are known indicators of motor function impairment in mice and have been used to monitor ALS progression in a mouse model (42). Hindpaws and forepaws are labeled separately with blue and red non-toxic ink, respectively. The subject is placed on a flat printing paper between two barriers two inches tall and four inches wide, ensuring straight, unidirectional walking, and allowed to walk for approximately 20 inches. The paper is scanned and reviewed in ImageJ. Four continuous pairs of hind pawprints establish the subject’s average hindlimb stride length: distance between the middle major knuckle of a pawprint and the same knuckle on that paw’s next print; stride width: distance between the midpoint of both stride length lines in the same gait cycle; and sway angle: angle between the direction the subject is traveling during a gait cycle, as determined by a line between the midpoints of the lines drawn between the middle major knuckles of each pair of hindlimb pawprints, and the direction each hindpaw’s final pawprint in that gait cycle as determined by a forward vector between its middle major knuckle and its furthest visible tarsal bone, either the calcaneus or talus. Degrees of parallel stepping are calculated by taking both hindpaw prints in a gait cycle and drawing a line from the middle distal of one to the other. Angles are formed where this line intersects the direction the mouse is traveling. Every pair of measurements thus sums to 180 degrees.

### Statistical Analysis

Differences across WT and SOD1 ALS mice were tested with mixed multivariate regression models using SAS v9.4 software. All other statistical analyses were performed in R (version 4.4.1; R Foundation, Vienna, Austria). Modeling considered the correlated observations within subjects with hierarchical linear modeling. Data were considered statistically significant where p < 0.05. Differences between BMS scores in left and right limbs were tested with two-tailed t-tests. Dendrograms of motor function decline were created via Ward clustering.

Correlations of motor function metrics’ rates of decline were established via the Pearson and Spearman methods; sex differences for motor function metrics were evaluated via Wilcoxon rank-sum tests and subjected to Benjamini-Hochberg correction.

---

**Supplemental Fig. 1.**
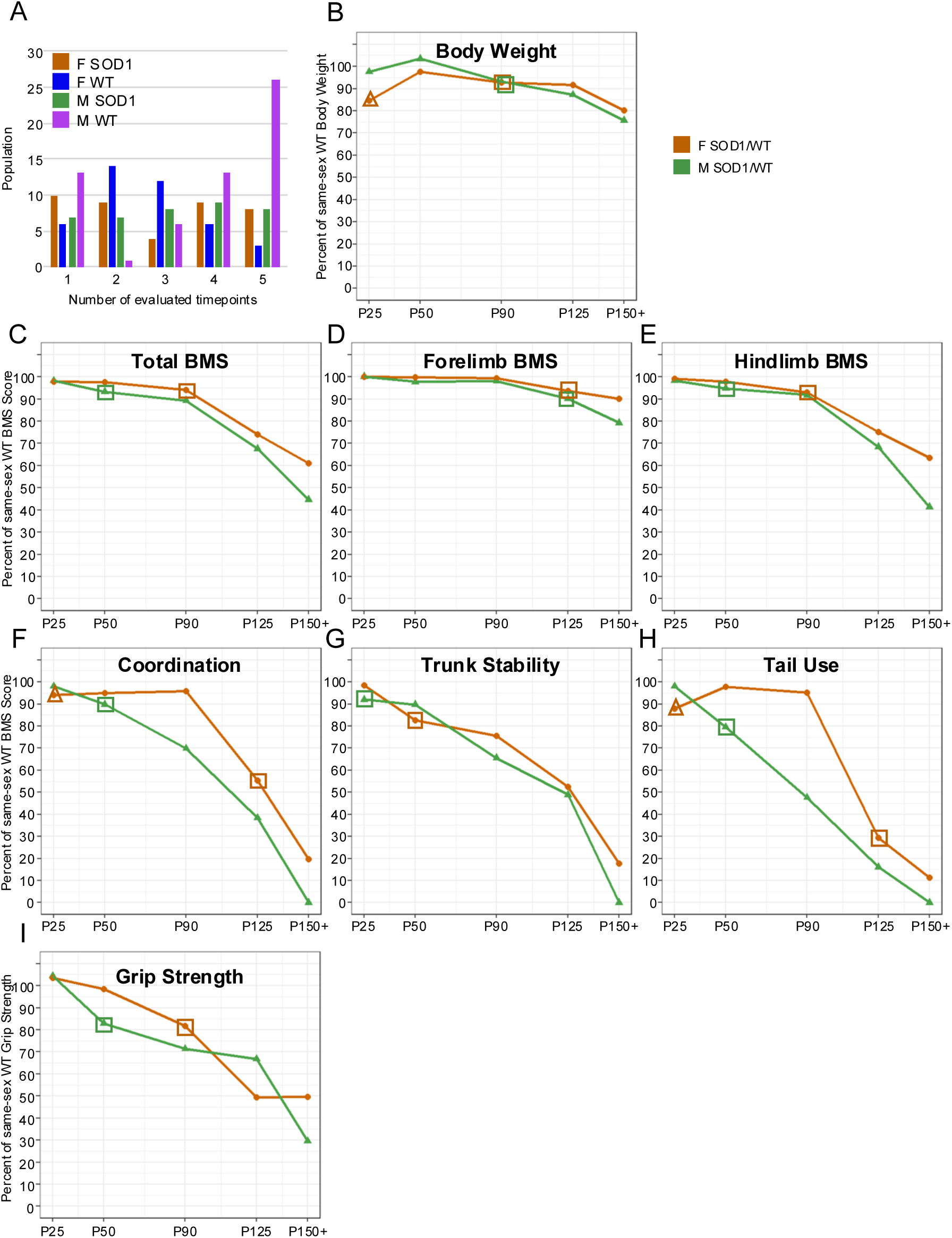
Divergence of motor function between SOD1 and WT occurs sooner than body weight divergence among males, and contemporaneously or later among females. A) The number of times mice of each sex-genotype population underwent evaluation. B) Expression of SOD1 body weight as a percentage of WT averages of the same sex and timepoint. C-I) Rate of BMS scores and grip strength as a percentage of WT averages of the same sex and timepoint. C-E: Overall, forelimb, and hindlimb BMS. F-H: The coordination, trunk stability, and tail use BMS criteria. I: Grip strength. Data points indicated by square outline denote ages where SOD1 average is reduced more than five percent below timepoint-matched WT average and subsequently continue to decrease. Data points indicated by a triangle outline denote features where the 25-day old SOD1 average is reduced more than five percent below 25-day old WT average, but subsequently increase to less than five percent difference.

**Supplemental Fig. 2.**
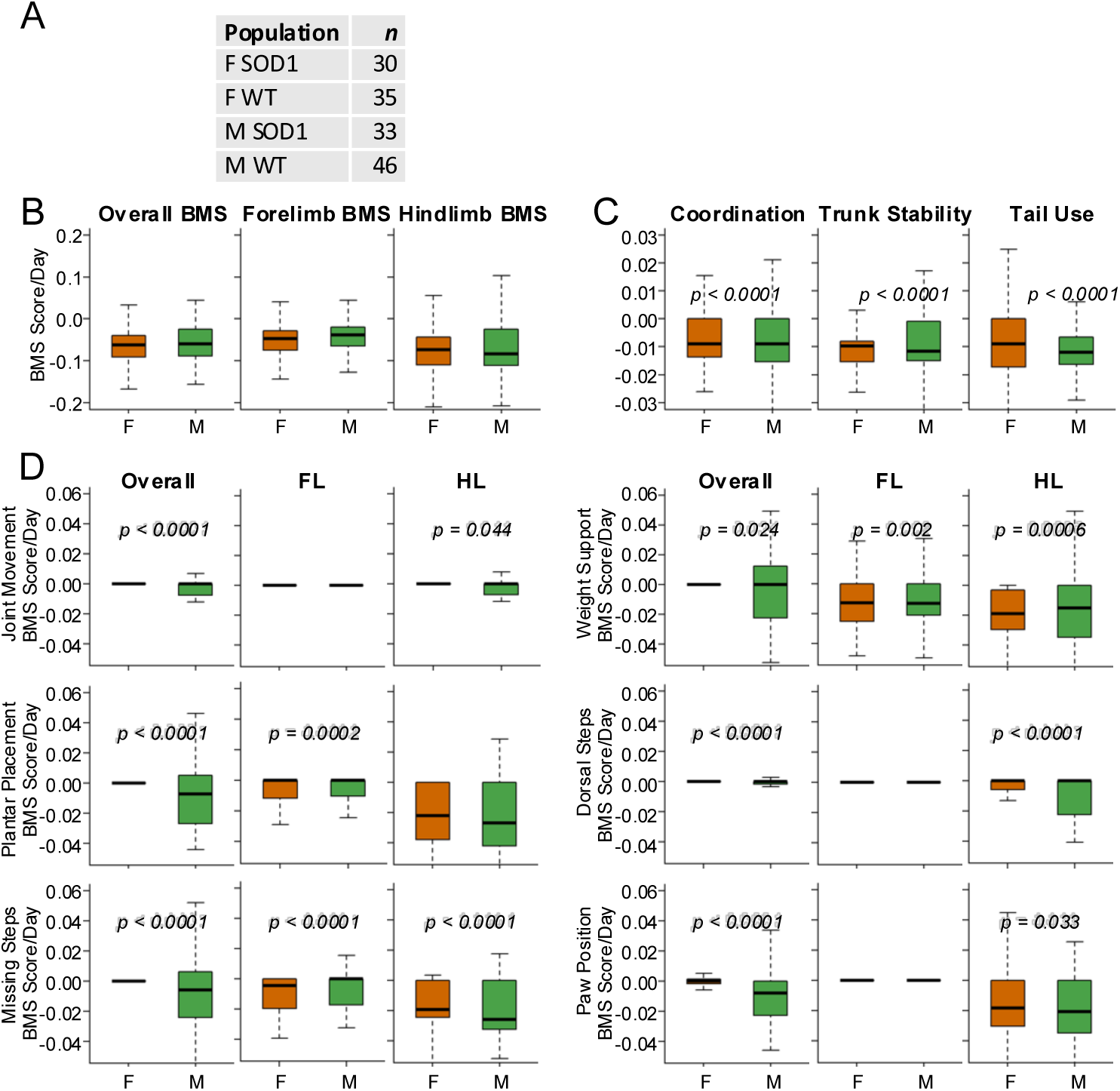
SOD1 transgenic BMS single-metric scores decline faster in males. Rates of change in BMS score difference from WT mice among SOD1 mice with at least two timepoints. Values are calculated from data shown in Fig 1 and 2. Wilcoxon rank-sum tests subject to Benjamini-Hochberg correction were used to determine p-values. A) Population of each sex-genotype combination. B) Overall, forelimb, and hindlimb BMS. C) Non-limb-specific BMS criteria. D) Limb-specific BMS criteria.

**Supplemental Fig. 3.**
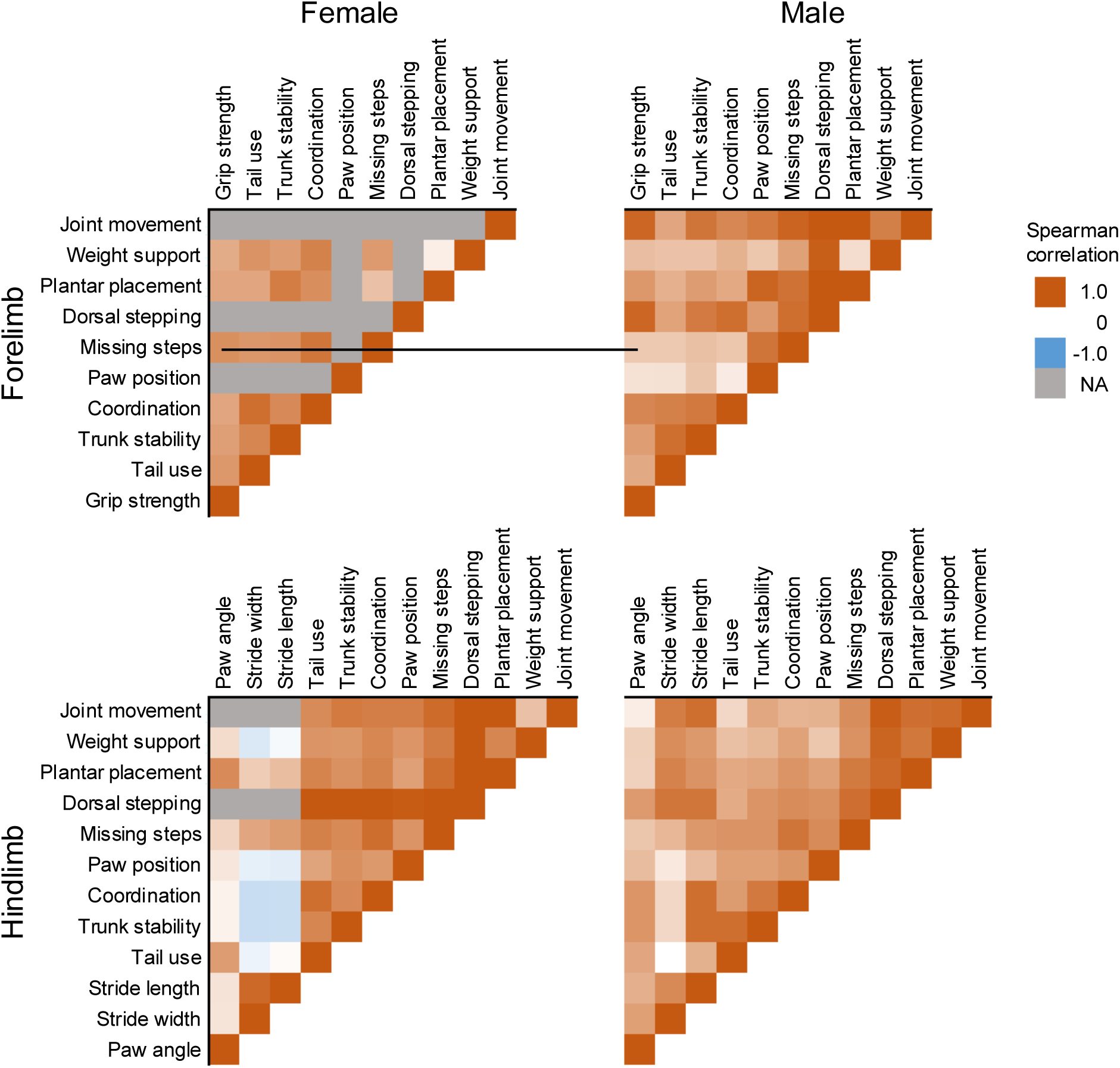
Correlations in decline between different motor behavior metrics differ between sexes. Average correlation of declining forelimb BMS and grip strength metrics with each other, as well as declining hindlimb BMS and pawprint test metrics with each other. Only SOD1 mice with multiple, longitudinal measurements in each assay were used to determine correlations. In metrics that did not change over time for any subject of that sex, correlations could not be calculated (colored grey, NA). Line indicates females exhibited a stronger correlation between forelimb missing steps and grip strength than males. N = x females and y males.

Supplemental Table 1. Data collected from SOD1 transgenic and WT mice at each timepoint. In the first sheet, data is organized by timepoint, which are displayed in order of increasing age from left to right. Rows represent mice, labeled by ID, sex, and genotype; mice of the same sex-genotype are displayed together. Each timepoint displays the following data over 70 columns (e.g. P25 stretches from Column B-BS). Col 1-2: mouse ID and body weight. Col 4-17: limb-specific BMS criteria for each limb (split into a HL and FL row for each subject as indicated by col 3; each criterion has both a left-limb (L) and right-limb (R) column). Col 18: sum of Paw Position scores for limbs indicated by the row at Lift Off and Initial Contact. Col 19-21: non-limb-specific BMS criteria. Col 22-30: sum BMS scores for the limbs indicated by column header (when multiple limbs’ BMS scores are summed, the contribution of each limb-specific criterion to the sum is the average of each limb’s score). Col 31-33/36-38/41-43: left-/right-/both-FL GS trial measurements, respectively. Col 34/39/44: average left-/right/both- GS measurements, respectively. Col 35/40/45: average left-/right-/both- GS measurements after normalization to body weight, respectively. Col 46-51: ink pawprint stride lengths. Col 52: average stride lengths. Col 53-57: stride widths. Col 58/59/60: average of stride widths from steps 1, 2, 3/steps 1.5, 2.5/all steps, respectively. Col 61-66: paw angles. Col 67/68/69: average right/left/all paw angles, respectively. Col 70: paw angle variance. Timepoints P90-P150 have 7 additional columns. Col 71-76: degrees of parallel stepping. Col 77: which HL has larger average degrees of parallel stepping. A “P150+” timepoint only displays body weight and BMS data for mice sacrificed after P150. Beneath each sex-genotype population are counts, averages, standard deviations, and confidence intervals for various displayed elements of that population. The second sheet contains a duplicate of the paw angle data, as well as original data on whether the subject leads gait with the left or right paw. The third sheet contains the ratio of average SOD1 mouse score over average WT mouse score for each sex-timepoint combination.

